# Characterizing the Effects of Filtration on Microbial Diversity, Reproducibility, and Genome Recovery from Wastewater

**DOI:** 10.64898/2026.11.17.767414

**Authors:** Nelson Ruth, Hajnalka Daligault, Tammie Martinez, Andy Hatch, Patrick Chain, Migun Shakya

## Abstract

Filtration is widely used in wastewater microbiome workflows, but how pore size, filtration tactics, and sampled fraction shape microbial profiles across sequencing approaches remains unclear. We compared direct and sequential filtration of wastewater through 3-, 0.45-, and 0.22-µm filters and analyzed filter-retained and corresponding filtrates using 16S rRNA gene and ITS amplicon sequencing, shotgun metagenomics, and metatranscriptomics. Filtrates generally yielded higher microbial alpha diversity but greater between-replicate variability, whereas filters produced more reproducible bacterial DNA profiles. Fraction-dependent differences were strongest for fungal and DNA viral communities, which separated between filters and filtrates, while bacterial DNA communities were largely shared across fractions. These patterns demonstrate that pore size does not act as a strict size cutoff in wastewater for bacteria. DNA viruses were consistently enriched in filtrates, which also yielded more viral genomes. In contrast, filters yielded more bacterial metagenome-assembled genomes, but overall bacterial genome recovery was limited. Bacterial and viral metatranscriptomic profiles were comparatively variable and did not support selection of a single preferred fraction for RNA-based recovery. Direct and sequential filtration also produced distinct profiles at the same terminal pore size, indicating that filtration history influences the recovered community. Together, these results suggest that wastewater processing workflow should therefore be selected according to the intended use case.

## Introduction

Wastewater sequencing has become a powerful approach for biosurveillance. While many wastewater surveillance networks target specific pathogens [1], broader characterization of microbial communities can be achieved through amplicon sequencing of marker genes such as 16S rRNA and ITS [2] or through untargeted metagenomic and metatranscriptomic sequencing. As wastewater surveillance continues to expand toward pathogen-agnostic detection, understanding how methodological choices shape microbial profiling has become increasingly important. As with all environmental sequencing workflows, wastewater sequencing is susceptible to biases introduced during sample collection, preprocessing, nucleic acid extraction, and sequencing. These biases can influence which organisms are ultimately detected [3]. For example, amplicon sequencing is affected by primer specificity, whereas shotgun sequencing can be influenced by sequencing depth and low nucleic acid yield. This makes systematic benchmarking essential for accurate interpretation of results.

Sample preprocessing is a particularly important source of bias. Comparative studies in microbial ecology, viromics, and metagenomics have demonstrated that choice of processing methods can alter estimates of microbial diversity, taxonomic composition, and genome recovery [3, 4]. Large interlaboratory comparisons of wastewater methods have shown that sample processing can substantially affect analytical sensitivity and reproducibility, emphasizing the need to benchmark workflows against their intended use cases [5]. These concerns are especially relevant to filtration. In marine systems, filtration studies have shown that microbial communities partition differently across filter pore sizes, with larger pore sizes often enriching particle-associated organisms while smaller pore-size membranes retain free-living microorganisms and extracellular nucleic acid [6, 7].

Filtration is among the most widely used steps for preprocessing wastewater samples [8, 9]. Wastewater contains a broad range of microbes with varying sizes including intact microbial cells, spores, virions, and microbial aggregates. Filtration is therefore likely to selectively partition specific bacterial, fungal, and viral groups between the filter and filtrate fractions. Characterizing these effects is necessary to interpret both detections and non-detections with confidence.

The impact of filtration may also differ depending on the sequencing approach. Amplicon sequencing can detect low-abundance bacterial and fungal taxa. In contrast, shotgun metagenomic and metatranscriptomic sequencing go beyond taxonomic profiling by recovering bacterial and viral genomes as well as capturing functional and transcriptional activity [10]. Despite these complementary strengths, few studies have evaluated how filtration influences microbial characterization across multiple sequencing approaches applied to the same wastewater samples.

Here, we evaluated how filtration pore size, filtration history, and sampled fraction influence microbial characterization across multiple sequencing approaches. Wastewater was processed by direct filtration or sequential filtration through 3-, 0.45-, and 0.22-µm membranes, and nucleic acids were extracted from both filter and corresponding filtrates. Bacterial and fungal communities were characterized by 16S rRNA gene and ITS amplicon sequencing, respectively, and microbial communities were evaluated by shotgun metagenomic and metatranscriptomic sequencing. We asked whether fraction-dependent patterns were consistent across microbial groups and sequencing approaches and whether the fractions that maximized taxonomic diversity also supported reproducible community profiling and genome recovery. By distinguishing the effects of pore size, prior filtration, and sampled fraction, this study provides a basis for tailoring wastewater filtration workflows to specific biosurveillance objectives.

## Results

### 1. Experimental design and sequencing overview

We designed an experiment to evaluate the effects of filtration pore size, sampled fraction (filter or filtrate), and sequencing approach for characterization of wastewater microbial communities (**Figure 1**). We conducted two filtration strategies: direct and sequential. In Direct Filtration (DF), we passed wastewater through a single filter with a pore size of 3 µm, 0.45 µm, and 0.22 µm. In Sequential Filtration (SF), we passed wastewater samples through filters of decreasing pore size, either from 3 to 0.45 µm, or 3 to 0.45 to 0.22 µm. DNA and RNA were extracted from both filter retentate and filtrate. For SF samples, only the final (smallest pore-size) filter and filtrate were used for nucleic acid extraction. DNA was used for 16S rRNA and ITS amplicon sequencing and shotgun metagenomic sequencing, and RNA was used for metatranscriptomic sequencing. The recovered DNA and RNA concentration from filters increased with decrease in pore sizes, but concentration of filtrate from samples were consistently lower for filtrate (**Supplementary Figure 1A**). Despite lower yield from filtrates, we still recovered similar amounts of sequences using both amplicon and shotgun-based methods (**Supplementary Figure 1B**).

**Figure 1:**
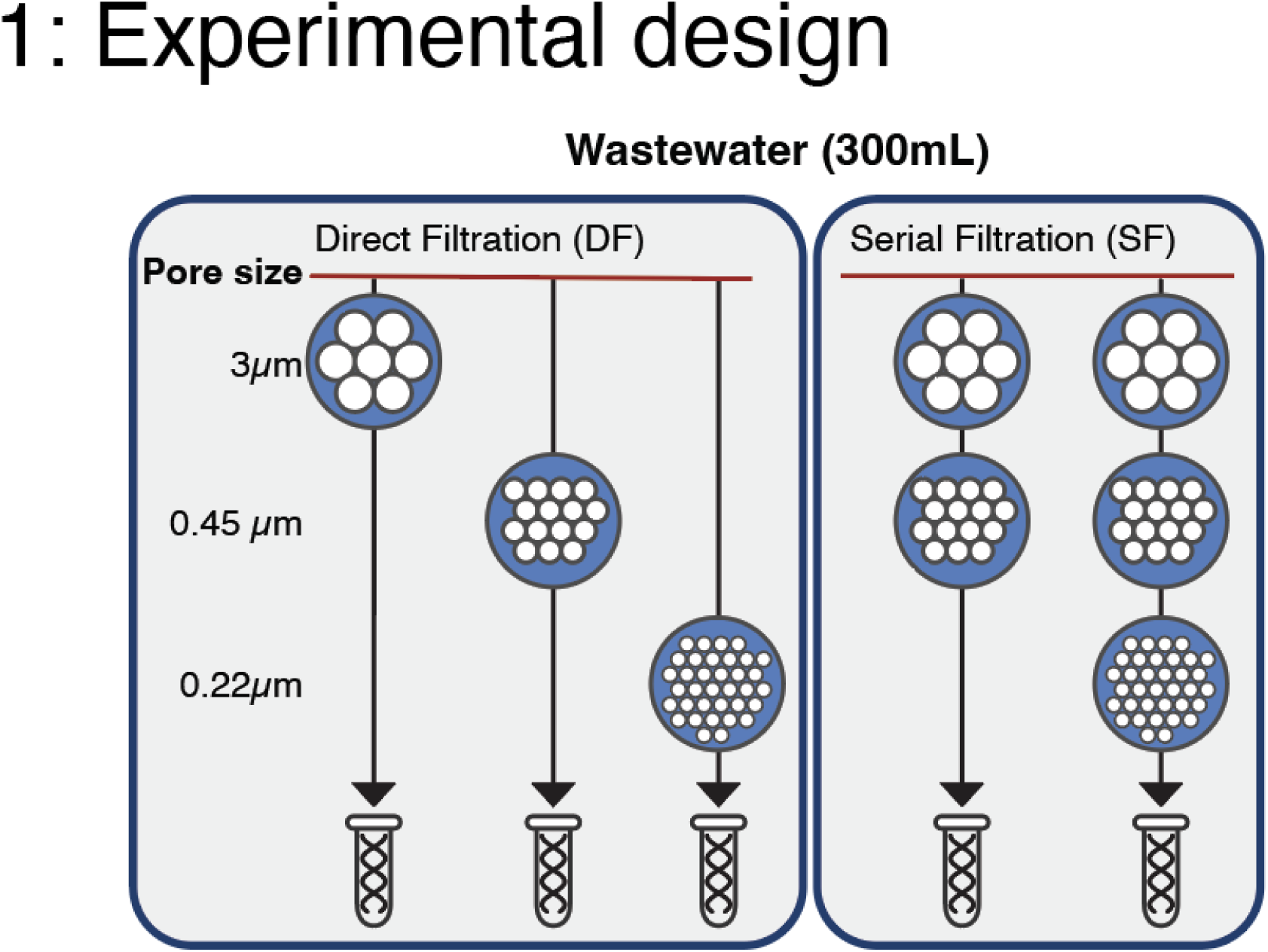
Overview of the experimental design. Wastewater samples were processed by either direct filtration (DF) through a single 3.0-, 0.45-, or 0.22-µm-pore-size membrane or sequential filtration (SF) through 3.0- and 0.45-µm membranes or through 3.0-, 0.45-, and 0.22-µm membranes. DNA and RNA were extracted separately from the filter-retained material and corresponding filtrates. For sequentially filtered samples, nucleic acids were extracted only from the terminal, smallest-pore-size filter and its corresponding filtrate. DNA was used for bacterial 16S rRNA and fungal ITS amplicon sequencing and for shotgun metagenomic sequencing. RNA was used for shotgun metatranscriptomic sequencing. metagenomic, and metatranscriptomic methods. Each filtration condition was processed in triplicate.

### 2. Filter and filtrate fractions differ in microbial diversity and reproducibility across sequencing approaches

Alpha diversity differed between filter and filtrate fractions, with filtrates showing higher but more variable diversity. In amplicon data, alpha diversity measured using the Effective Number of Species (ENS; Hill number) was higher in filtrates than filter for fungal and bacterial communities (**Figure 2A and 2B**), with bacterial ENS being highest in SF 0.22 µm filtrates, suggesting that the removal of larger particles results in improved recovery of broader diversity. Furthermore, filtrates were less reproducible than filters, with a mean inter-replicate coefficient of variation (CV) of 38.6% compared with 15.7% for filters.

**Figure 2.**
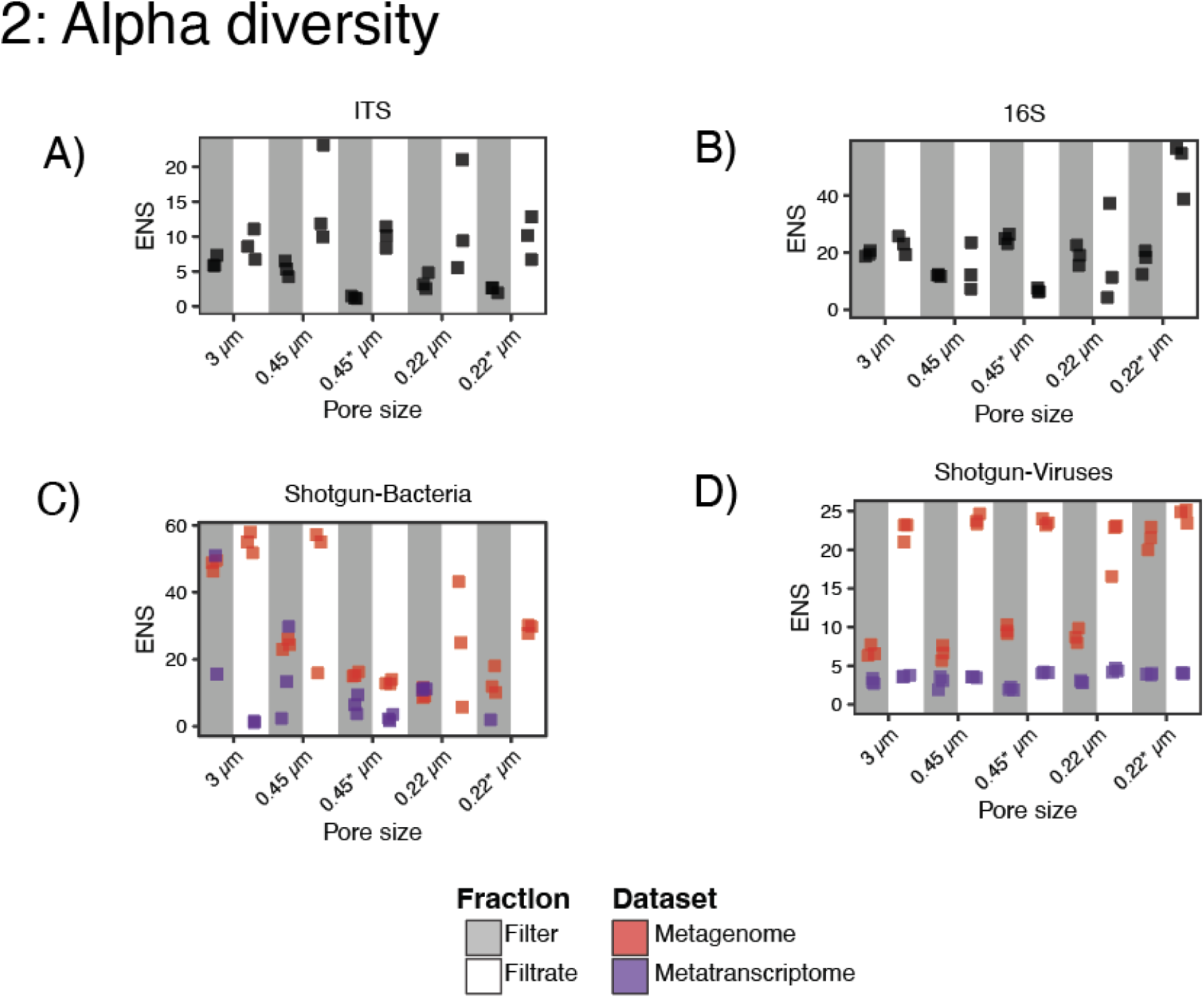
Alpha diversity across filtration fractions and sequencing approaches. Alpha diversity is reported as the effective number of species (ENS), calculated as the Hill number of order (q=1). ENS values are shown for microbial and viral communities recovered from filter-retained material and corresponding filtrates following direct (DF) or sequential filtration (SF) using 3.0-, 0.45-, and 0.22-µm-pore-size membranes. Panels A and B show fungal and bacterial communities profiled by ITS and 16S rRNA gene amplicon sequencing, respectively. Panels C and D show bacterial and viral communities, profiled by shotgun metagenomic and metatranscriptomic sequencing.

To assess diversity in the shotgun datasets, we used taxonomic classifications derived from metagenomic and metatranscriptomic reads. Analyses were limited to bacterial and viral communities because fungal and archaeal reads were sparse in these datasets. In the metagenomic data, bacterial ENS was consistently higher in filtrates than in filters (**Figure 2C**). However, bacterial profiles were more reproducible in filters, with a mean CV of 12.4%, compared with 29.2% in filtrates. DNA viral ENS was also higher in filtrates. The SF 0.22-µm filter was an exception, with a viral ENS of 21.5, comparable to values observed in the filtrates. The bacterial metatranscriptomic profiles showed a different pattern from the DNA-based profiles. Variability was high in both fractions and was greater in filters (mean CV = 53.0%) than in filtrates (mean CV = 31.9%) (**Figure 2D**). Thus, the greater reproducibility of filter-retained bacterial profiles observed in the metagenomic data did not extend to the metatranscriptomic data.

### 3. Filter–filtrate differences in community composition are strongest for fungi and DNA viruses

To determine whether fraction type and filtration strategy influenced microbial community composition, we performed principal coordinate analysis (PCoA) based on Bray–Curtis dissimilarities (**Figure 3**). The degree of separation between filter and filtrate communities varied based on taxonomic group and sequencing approach. Separation was clearest in fungal communities, where filters and filtrates formed two largely distinct clusters, each containing both SF and DF samples (**Figure 3A**) (PERMANOVA: pseudo-*F* = 12.78, R^2^ = 0.313, adjusted *P* = 0.0015). However, variability among replicate community profiles also differed between fractions (PERMDISP: adjusted *P* = 0.0012), indicating that it also contributed to observed separation. In contrast, 16S rRNA amplicon datasets showed no clear separation between filter and filtrate communities, although SF 0.22 µm filter samples formed a distinct cluster (**Figure 3B**). A PERMANOVA comparing their bacterial communities with those from all DF filters combined supported this pattern (pseudo-*F* = 21.14, R^2^ = 0.679, adjusted *P* = 0.0044).

**Figure 3.**
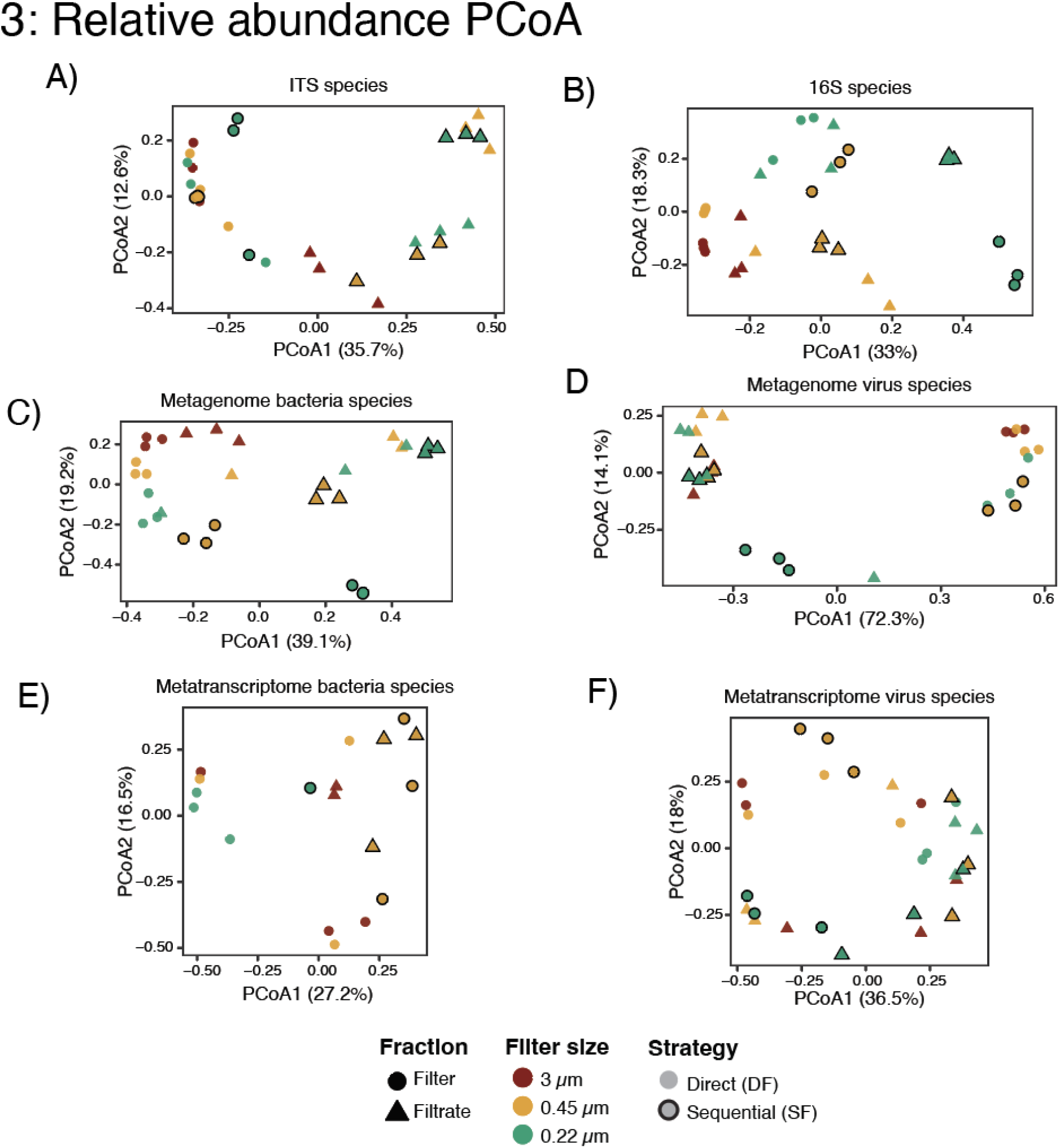
Principal coordinates analysis of wastewater microbial and viral communitie across filtration strategies. Principal coordinates analysis (PCoA) based on Bray–Curtis dissimilarities visualize patterns of similarity and dissimilarity among communities recovered from filter and corresponding filtrates produced by direct (DF) and sequential filtration (SF). Panels A and B show fungal and bacterial communities profiled by ITS and 16S rRNA gene amplicon sequencing, respectively. Panels C and D show bacterial and DNA viral communities, respectively, profiled by shotgun metagenomic sequencing. Panels E and F show bacterial and RNA viral communities, respectively, profiled by shotgun metatranscriptomic sequencing. Each point represents one independent filtration replicate (n = 3 per condition). Point shape denotes the sampled fraction (circle = filter; triangle = filtrate), color denotes the terminal membrane pore size (red, 3.0 µm; orange, 0.45 µm; green, 0.22 µm), and marker outline denotes the filtration strategy (no outline = DF; dark outline, SF).

Bacterial species profiles from metagenomes partly mirrored the 16S rRNA results, with no clear separation between filters and filtrates (**Figure 3C**). DNA viral communities showed separation similar to fungal communities with filters and filtrate forming distinct clusters regardless of pore size (pseudo-F = 36.41, (R^2) = 0.543, adjusted P = 0.0015) (**Figure 3D**). The SF 0.22-µm filter samples also appeared closer to filtrates in the ordination, consistent with their similar viral ENS values (**Figure 2D**). Metatranscriptomic-based profiles showed weak structures, with no clear clustering for both bacterial and viral fraction (**Figure 3E, 3F**). Together, these results indicate that fungal and DNA viral communities are strongly influenced by whether filters or filtrates are sampled. Bacterial composition was influenced less by fraction type and more by specific conditions, particularly SF 0.22 µm filtration. Metatranscriptomic profiles were less reproducible than DNA-based profiles, consistent with their high inter-replicate variability in alpha diversity.

### 4. Taxonomic partitioning between filters and filtrates differs among bacteria, fungi, and viruses

Beyond the differences in community composition, taxon-level comparison revealed that bacteria, fungi, and viruses partitioned differently between filters and filtrates. We assessed this by identifying taxa enriched in, or exclusive to, each fraction. Here, “enriched” refers to a higher relative sequence abundance in one fraction and does not imply greater absolute recovery per unit volume. Bacterial communities showed the weakest partitioning. Based on the 16S rRNA amplicon data, most bacterial classes were detected in both filters and filtrates (**Supplementary Figure 4A**). However, class *Saccharimonadia*, a class of bacteria consisting of ultra small bacteria, were enriched in all except 0.22 µm SF filtrates (**Supplementary Figure 2A**), consistent with its ability to pass through conventional membrane filters [11, 12]. Bacterial communities inferred from metagenomes showed similar trend, with most classes shared between fractions (**Supplementary Figure 4B**). However, *Saccharimonadia* was enriched only in the DF 3 and 0.45 µm filtrates (**Supplementary Figure 2C**). Metatranscriptomic profiles were sparser, partly because of lower sequencing depth. Even so, many bacterial classes were detected only on filters (**Figure 4B**), and no class was enriched across all filtrates. Thus, DNA from filter-passing bacteria persists in filtrates, but transcriptionally active bacteria were associated primarily with biomass retained on filters.

**Figure 4.**
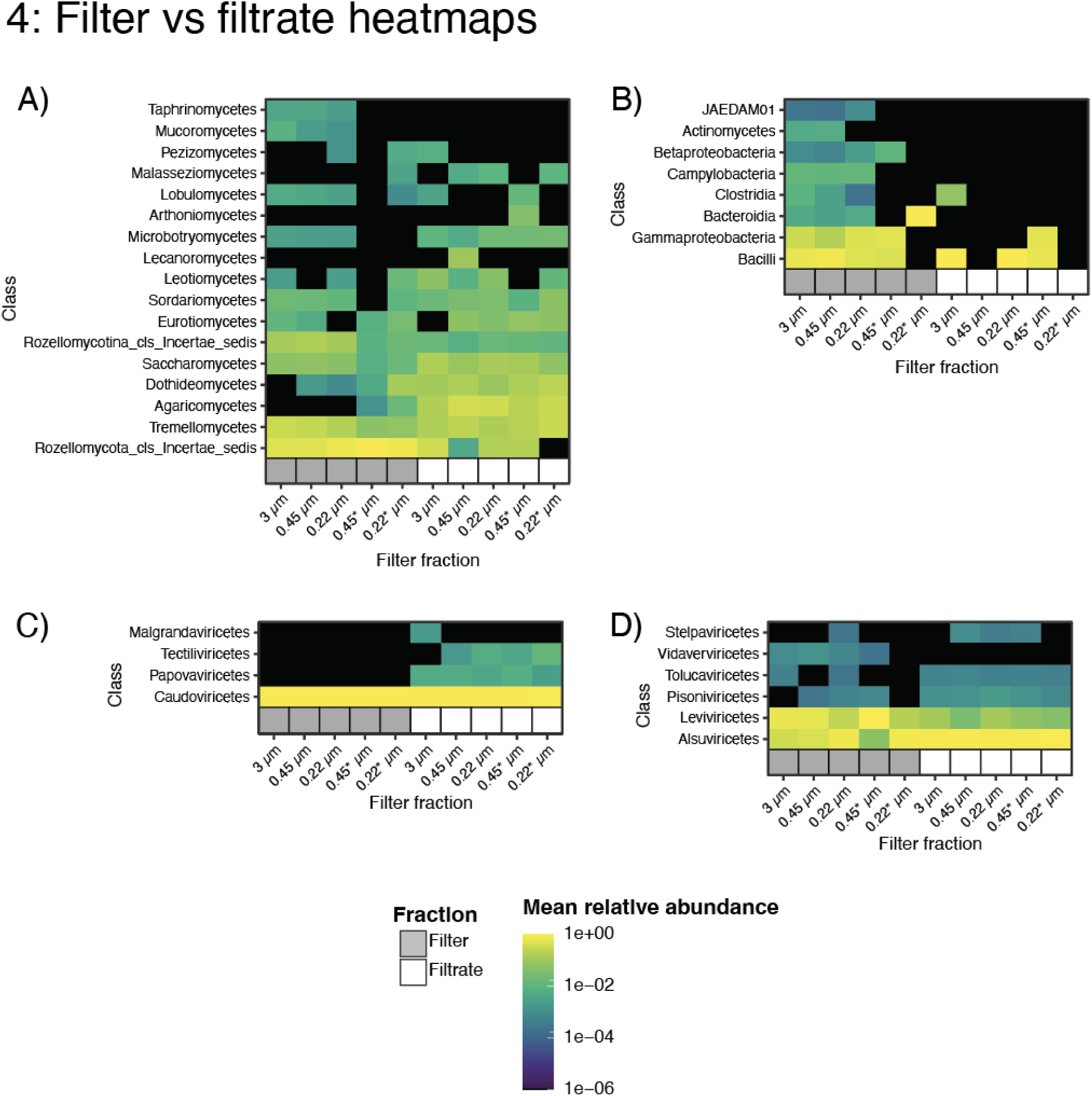
Relative abundance of microbial and viral taxonomic classes across wastewate filtration fractions. Heatmaps show the mean relative sequence abundance, expressed as a proportion, of taxonomic classes detected in filter and corresponding filtrates produced by direct (DF) and sequential filtration (SF) using terminal membrane pore sizes of 3.0, 0.45, and 0.22 µm. Panels show (A) fungal classes profiled by ITS amplicon sequencing, (B) Bacterial classes profiled from metatranscriptomes (C) DNA viral communities profiled from metagenomes, and (D) RNA viral classes profiled by metatranscriptomes. Each column represents the mean relative abundance across three independent replicates. Only taxonomic classes with an overall mean relative abundance greater than 0.001 across all samples are shown. Blank cells indicate a relative abundance of zero. Relative abundance values were obtained from GOTTCHA2 taxonomic profiles.

Fungal communities showed stronger, filtrate-based partitioning. Several fungal classes including *Agaricomycetes,* were enriched in filtrates regardless of the pore size (**Supplementary Figure 2B**). This partitioning was further highlighted by the exclusive recovery of specific fungal class within a single filtration (e.g. *Arthoniomycetes* only in 0.45 µm filtrate of SF) (**Figure 4A**). At the species level, one fungal species (Uncultured GS08_sp) was detected across all methods and fractions (**Supplementary Figure 3A**). While the DF 0.45 µm filtrate recovered the most fungal species (n = 35), the DF 0.22 µm filtrate captured the most (n=15) unique species. This filtrate-exclusive recovery demonstrates that fungal diversity persists even at fine scales, likely manifesting as small particulates, structural fragments, or extracellular forms capable of passing through 0.22 µm membranes.

Viral communities showed the most consistent partitioning. Nearly all DNA virus classes were enriched in filtrates across both DF and SF fractions. Tailless DNA viral classes such as *Papovaviricetes* and *Tectiliviricetes* were enriched in most filtrate fractions regardless of pore size (**Figure 4C**). At the species level, DNA viral species recovery differed from bacteria in exhibiting broader overlap of species across filtration strategies, with a high number of shared viral species consistently recovered from filtrates (**Supplementary Figure 3C**). In contrast, RNA virus enrichment patterns varied across fractions and filtration conditions. *Vidaverviricetes*, a class containing bacterial-infecting RNA viruses, was enriched on filters rather than in filtrates in all DF and in the SF 0.45 µm fraction (**Figure 4D**). This contrasting behavior suggests that RNA viral recovery may be more influenced by attachment to host cells, biofilm particles, or extracellular aggregates than DNA viruses, leading to retention within filters despite their small sizes.

### 5. Bacterial and viral genomes partition differently between filters and filtrates

Recovery of bacterial metagenome-assembled genomes (MAGs) was limited across all samples (peaking at a maximum of 5 MAGs in one sample), reflecting insufficient sequencing depth needed to recover MAGs. Despite this limitation, genome recovery differed consistently between fractions. Filters yielded more bacterial MAGs than their corresponding filtrate, which recovered (0-3) genomes. Notably, the SF 0.22 µm filtrate yielded zero bacterial MAGs (**Figure 5A**). In contrast, filtrates yielded more and higher-quality DNA viral genomes than their corresponding filters (**Figure 5B**). However, this predictable partitioning did not extend to the metatranscriptomic data. Recovery of high-quality RNA viral genomes was highly variable, and neither filters nor filtrates consistently produced higher-quality genomes. This suggests that while DNA-based viral genomes partition predictably into the filterable fraction, the recovery of RNA virus was more variable.

**Figure 5.**
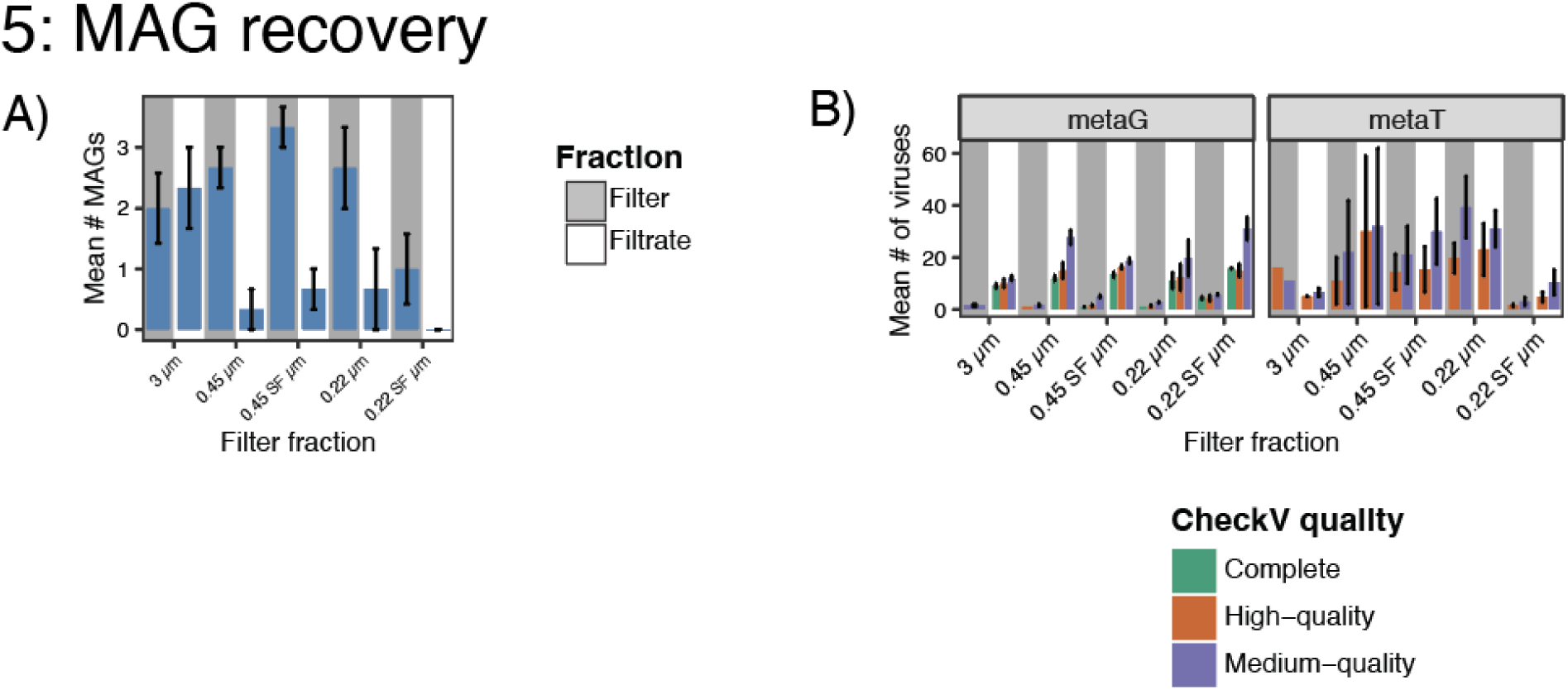
Recovery of bacterial metagenome-assembled genomes and viral genomes across wastewater filtration fractions. (A) Average number of bacterial metagenome-assembled genomes (MAGs) meeting medium-quality or higher criteria recovered from shotgun metagenomic assemblies of filter-retained material and corresponding filtrates. (B) Average number of DNA and RNA viral genomes recovered from metagenome and metatranscriptome. Bars represent the mean across three independent filtration replicates, and error bars represent ±1 standard error of the mean (SEM).

## Discussion

This study demonstrates that filtration is not a neutral preprocessing step in wastewater sequencing. Filter retentates and filtrates derived from the same sample produced different microbial community, and the magnitude of these differences depended on the organismal group, nucleic acid type, and sequencing approach. Fraction-dependent separation was strongest for fungal and DNA viral communities, whereas bacterial DNA profiles were broadly shared between filters and filtrates. Filtrates generally recovered higher number of species but also showed greater variability between replicates, while filters showed more consistent bacterial DNA profiles and higher bacterial MAG recovery. DNA viral genomes recovery showed the opposite pattern, and recovered genomes were higher in filtrates. Metatranscriptomic profiles were comparatively more variable in both fractions. Collectively, these results argue against a universally optimal wastewater reaction and wastewater processing workflow selection should be based on whether the objective is reproducible bacterial community profiling, fungal or viral detection, or recovery of bacterial versus viral genomes.

The pattern from the SF 0.22-µm samples further shows that pore size alone does not effect the recovered fraction. These samples had first passed through 3-and 0.45-µm membranes and were repeatedly distinct from samples collected by direct filtration. Upstream filters can remove aggregates, particle-associated organisms, and extracellular material before filtering through final pore size. Volume- and biomass-dependent filtration biases have altered both alpha and beta diversity in aquatic microbial community studies [13], and broad protocol comparisons have found size fractionation to be a major determinant of recovered community structure [14]. Thus, a 0.22-µm membrane used after prefiltration does not sample the same analyte pool as a 0.22-µm membrane applied directly to wastewater. Thus, Filtration history should be accounted for when interpreting results.

Bacteria showed the weakest overall partitioning between filters and filtrates. Most bacterial classes were detected in both fractions by 16S rRNA amplicon sequencing and metagenomics, consistent with contributions from intact cells spanning a range of sizes, particle-associated and free-living populations, and extracellular DNA. *Saccharimonadia* provided a biologically plausible exception because members of the candidate phyla are usually defined by its ultrasmall cell size. Bacteria from related CPR lineages have been recovered after passage through approximately 0.2-µm filters [15] and cultured *Saccharibacteria* have been isolated by passing oral samples through 0.2-µm membranes before coculture with bacterial hosts [16]. Their enrichment in several wastewater filtrates is therefore consistent with filter passage by small cells, although we cannot distinguish these from extracellular or cell-associated DNA in the present study. The failure to observe enrichment in every filtrate also suggests that upstream retention, host or particle association, or sampling variability modifies this size-based tendency.

Fungal communities exhibited clearer filter–filtrate separation than bacterial communities, demonstrating that fungal surveillance is particularly sensitive to the sampled fraction. The recovery of numerous fungal taxa, including filtrate-exclusive taxa, after passage through 0.22-µm membranes is unlikely to be explained solely by intact vegetative fungal cells. Small spores, hyphal or cellular fragments, extracellular DNA, and deformation or breakthrough during filtration are all plausible contributors. ITS amplicon data cannot distinguish among these possibilities. Nevertheless, the result is operationally important: a wastewater filtrate should not be considered devoid of fungal signal, and analysis of filter material alone may miss relevant fungal targets.

DNA viruses showed the most consistent enrichment in filtrates, which also yielded more viral genomes with higher CheckV-estimated quality. This pattern is compatible with passage of viral particles through the membranes and also supports the use of filtration in wastewater viromics [17]. Importantly, small size of viruses alone cannot explain partitioning. Associations of some viruses with their host may offer a plausible explanation for the recovery of viral groups on filters and for the enrichment of *Vidaverviricetes* in several filter fractions. The contrast between DNA viral enrichment in filtrates and variable RNA viral community pattern is likely the combined outcome of viral ecology, particle association, nucleic acid stability, and distinct DNA and RNA protocols used for processing them.

Amplicon and shotgun sequencing reproduced some broad bacterial patterns, particularly the extensive overlap between filter and filtrate communities, but they were not interchangeable at finer taxonomic resolution. Past comparisons using environmental samples have shown that 16S rRNA amplicon sequencing can recover taxa missed by shallow shotgun sequencing [10]. Both approaches also contain taxon- and protocol-specific measurement biases [18]. In the present study, the comparatively stable DNA-based bacterial patterns suggest that filters may be preferable when reproducible community comparisons or genome reconstruction are primary goals. Conversely, filtrates were more efficient for DNA viral diversity and genome recovery. Fungal profiling may benefit from retaining and analyzing both fractions, whereas the variable metatranscriptomic results do not yet support selecting a single fraction for RNA virus recovery.

The low number of bacterial MAGs limits genome-level inference but reinforces the effect of fractionation. Filters contained more biomass and yielded more bacterial MAGs, whereas the corresponding filtrates—especially the sequentially filtered 0.22-µm fraction—yielded few or none. MAG recovery depends on sequencing depth, organism abundance, community complexity, and taxonomic relatedness, and low recovery cannot be attributed to filtration alone [19]. Accordingly, these data support a qualitative difference between fractions but do not establish the quantitative efficiency of bacterial genome recovery. Deeper sequencing is required to determine whether filtrate-associated bacteria can be reconstructed reliably.

Several limitations define the scope of our work. First, the absence of spike-ins prevents calculation of efficiency and taxon-specific bias from being measured against a known ground truth. Second, the analyses are based primarily on relative sequence abundance. An apparent enrichment can result either from increased recovery of a target or from depletion of other community members. Third, the study did not distinguish intact cells or virions from free or particle-associated nucleic acids. Fourth, the 300-mL input volume may have increased membrane loading and clogging; because filtration bias can depend on processed volume and biomass [13], the results should not be extrapolated directly to workflows using substantially different volumes. Finally, broader validation across wastewater sources, seasons, and physicochemical factors is needed before a single protocol can be recommended.

In summary, pore size, filtration history, sampled fraction, and sequencing strategy jointly determine the microbial signal recovered from wastewater. Filters provided the most reproducible bacterial DNA profiles and favored bacterial MAG recovery, whereas filtrates captured greater but more variable diversity and favored DNA viral detection and viral genome recovery. Fungal and RNA viral signals did not conform to a simple size-exclusion model. Despite these limitations, benchmarking studies like ours remain important as wastewater surveillance continues to expand into large-scale public health applications. The results presented here demonstrate that methodological choices influence observed microbial and viral recovery patterns, emphasizing the need for standardized benchmarking frameworks capable of evaluating method-specific biases for specific surveillance goals.

## Methods

### Wastewater sampling and sample preparation

Twenty-four-hour composite wastewater samples were collected over two consecutive days (February 7–8, 2024) from the Los Alamos National Laboratory wastewater treatment plant. A total volume of 6.5 L was pooled for processing. To remove extracellular DNA (eDNA), the pooled wastewater was treated with 1,000 U (0.5 mL) of DNase I (M0303S, NEB) at room temperature overnight. Following treatment, samples were centrifuged at 3,500 × g for 10 min to remove large particulates and bulk solids. Approximately 5.85 L of clarified supernatant was recovered for subsequent fractionation experiments.

### Experimental filtration workflows and biomass fractionation

All experiments were performed in triplicate (n=3) using 300 mL aliquots of clarified wastewater. Filtration method was done in two broad approaches: Direct Filtration (DF) and sequential filtration (SF). In DF Samples were passed through individual MF-Millipore mixed cellulose ester (MCE) membranes with pore sizes of 3.0 µm (DF1; SSWP04700), 0.45 µm (DF2; HAWG04700), or 0.22 µm (DF3; GSWP04700). In SF wastewater was first passed through a 3.0 µm membrane, and the resulting filtrate was subsequently filtered through a 0.45 µm membrane (SF1). In a secondary sequential regime (SF2), the filtrate from the 0.45 µm step was further passed through a 0.22 µm membrane to assess the effect of multi-stage fractionation on biomass recovery (**Figure 1**).

### Nucleic acid extraction and quality control

Filters and filtrate fractions were processed immediately following each filtration experiment. Each membrane was bisected; one half was utilized for RNA isolation (RNeasy PowerWater Kit, Qiagen) and the remaining half for DNA isolation (DNeasy PowerWater Kit, Qiagen). Filtrate fractions (15 mL/tube) were concentrated using Amicon Ultra Centrifugal Filters (10 kDa MWCO; Millipore) at 5,000 × g for 20 min. The resulting concentrates were split equally for DNA and RNA extraction using the aforementioned kits. Total DNA and RNA concentrations were quantified using Qubit High Sensitivity assays (Thermo Fisher Scientific)

### Amplicon sequencing

For amplicon-based taxonomic profiling, the 16S rRNA gene was amplified using primers F341 (5′-CCTACGGGNBGCASCAG-3′) and R806 (5′-GGACTACHVGGGTWTCTAAT-3′). The 16S library was prepared using the Quick-16S™ NGS Library Prep Kit (Zymo Research, Irvine, CA).

Fungal communities were profiled by amplifying the ITS3–ITS4 region using primers ITS3 (5′-GCATCGATGAAGAACGCAGC-3′) and ITS4 (5′-TCCTCCGCTTATTGATATGC-3′). The first PCR used KAPA HiFi HotStart ReadyMix (Kapa Biosystems, KK2602) with an initial denaturation at 95°C for 3 min; 20 cycles of 95°C for 30 s, 55°C for 30 s, and 72°C for 30 s; and a final extension at 72°C for 5 min. A second PCR added Illumina sequencing adapters and unique indexes using the Nextera XT Index Kit v2 (Illumina, FC-131-2001 through FC-131-2004) and KAPA HiFi HotStart ReadyMix. This reaction used an initial denaturation at 95°C for 3 min; eight cycles of 95°C for 30 s, 55°C for 30 s, and 72°C for 30 s; and a final extension at 72°C for 5 min. Amplicons were purified with AMPure XP beads (Beckman Coulter, A63881).

A no-template control and positive control were included for ITS library preparation; the no-template control showed no band at the expected amplicon size and was sequenced for confirmation. For 16S library preparation, ZymoBIOMICS® Microbial Community DNA Standard (Zymo Research) was used as a positive control, and blank extraction and library-preparation controls were included to assess laboratory contamination. Libraries were quantified using the Qubit dsDNA HS Assay (Thermo Fisher Scientific, Q32854), assessed for size using the Agilent High Sensitivity DNA Kit, and accurately quantified using the KAPA Library Quantification Kit–Illumina/Universal (Kapa Biosystems, KK4824). Indexed libraries were pooled and sequenced on an Illumina NextSeq 2000 using a P1 Reagent Kit.

### Metatranscriptomic sequencing

The concentration of the total RNA was obtained using the Qubit RNA HS Assay and quality was determined using Agilent Bioanalyzer RNA Pico Assay. Ribosomal RNA was depleted from total RNA using the Illumina Ribo-Zero Plus rRNA Depletion Kit (Illumina, Cat. #20037135). RNA was converted to cDNA and adapters and indexes were added onto the ends of the fragments to generate Illumina libraries that can be sequenced on an Illumina sequencer. Illumina libraries were eluted in DNA Elution Buffer (Zymo Research, Cat. #D3004-4-10). The concentration and average size of the libraries was obtained using the Tapestation DNA 5000 Assay Reagents and Screen Tapes (Agilent, Cat. #5067-5589 and 5067-5588). Libraries were normalized and pooled based on the Tapestation concentrations. An accurate library quantification of the pool was determined using the Library Quantification Kit – Illumina/Universal Kit (KAPA Biosystems, KK4824). The library pool was sequenced to generate paired-end 151 bp reads using the NextSeq 2000 P3 Reagent Kit (300 cycles) (Illumina).

### Metagenomic sequencing

Metagenomic Illumina libraries were prepared using NEBNext Ultra DNA Library Preparation Kit (New England Biolabs, Cat. #E7370S) according to the manufacturer’s protocol. DNA was fragmented using a Covaris E220, the ends blunted, and adapters and indices were added onto the ends of the fragments prior to PCR enrichment. For samples with inputs less than 10 ng of DNA, the number of PCR cycles was increased from the manufacturer’s protocol of 10 to 15 cycles. AMPure XP beads were used to clean the Illumina libraries (Beckman Coulter, Cat. #A63881) prior to elution in DNA Elution Buffer (Zymo Research, Cat. #D3004-4-10). Library concentrations were obtained using the Qubit dsDNA HS Assay (ThermoFisher Scientific, Cat. #Q32854). The average size of the library was determined by the Agilent High Sensitivity DNA Kit (Agilent, Cat. #5067-4626). An accurate library quantification was determined using the Library Quantification Kit – Illumina/Universal Kit (KAPA Biosystems, Cat. #KK4824). Prepared libraries were sequenced on the Illumina NextSeq 500 to generate paired-end 151 bp reads.

### Bioinformatic processing of amplicon datasets

Taxonomic profiling of the prokaryotic (16S rRNA) and fungal (ITS) communities was performed using the QIIME [v2023.5] [20] integrated within the EDGE Bioinformatics platform [v2.4.1] [21]. Specifically, we used DADA2 pipeline, which performs quality control, denoises quality reads, and filters out chimeras using the parameters: *Trim 5*′ *end Forward = 20, Trim 5*′ *end Reverse = 20, Truncation Len Forward = 0, Truncation Len Reverse = 0, SE Trim Length = 20, SE Truncation Len = 0*. After quality control and filtering, reads were clustered into Amplicon Sequence Variants (ASVs). Taxonomic assignment for ASVs was conducted using the GreenGenes database [v2023.5] [22] for 16S, while fungal ITS sequences were classified using the UNITE database [v2022_10] [23]. For amplicon datasets, alpha diversity was expressed as coverage-standardized effective number of species (ENS; Hill number, (q=1)).

Beta diversity was visualized by PCoA of Bray–Curtis dissimilarities calculated from coverage-standardized species-level abundance matrices. Each dataset was analyzed separately using PERMANOVA, accounting for matched filter–filtrate pairs and restricting permutations to preserve the dependence between successive stages of each sequential-filtration run. A total of 9,999 permutations was requested, with complete enumeration when fewer valid permutations were available. Pseudo-F statistics, R², P values, and actual permutation counts were reported, with Benjamini–Hochberg adjustment within comparison families. Homogeneity of multivariate dispersion was assessed using PERMDISP with spatial medians, small-sample bias correction, square-root Bray–Curtis distances, and corresponding permutation restrictions.

### Read-based taxonomic profiling of metagenomes and metatranscriptomes

Raw metagenomic and metatranscriptomic reads were subjected to quality control and adapter trimming using the FaQCs [v2.12] [24] where reads were filtered to remove sequencing adapters, low-quality bases (Phred score < 20), and short fragments (< 50 bp). Taxonomic assignment for the quality-filtered shotgun reads was performed using GOTTCHA2 [v2.2.3][25].

For all downstream analyses (e.g. Hill diversity calculations, Bray-Curtis), taxonomic classifications were restricted to species level to maintain high-resolution taxonomic precision.

### Enrichment and depletion analyses

QIIME2 outputs were imported into R [v4.5.3] and organized into phyloseq [v1.54.2] objects using ‘otu_table()’, ‘sample_data()’, ‘tax_table()’, and ‘phyloseq()’. Following taxonomic filtering, 16S and ITS datasets were agglomerated to species level using ‘tax_glom()’. For metagenomic and metatranscriptomic datasets, bacterial and viral species-level read-count matrices were generated from GOTTCHA2 outputs without a fixed minimum read-count threshold. Across all datasets, alpha diversity was estimated using coverage-based rarefaction/extrapolation in iNEXT [v3.0.2] and expressed as the effective number of species (ENS; Hill number of order (q=1)), equivalent to the exponential of Shannon entropy. Mean ENS ± SD was summarized by dataset and filter/filtrate fraction. Technical reproducibility was assessed using replicate CVs (100 × SD/mean), calculated within each filtration condition and averaged separately for each dataset and fraction.

Taxonomic composition heatmaps were generated at the class level for each dataset. For the QIIME-derived 16S and ITS datasets, taxa were agglomerated to the class level using ‘tax_glom()’ and converted to within-sample relative abundances; the ITS analysis was restricted to fungi. For the GOTTCHA2-derived metagenomic and metatranscriptomic datasets, relative abundances were summed by class within each sample and renormalized separately within the retained bacterial and viral subsets. Class-level relative abundances were then averaged across replicates within each filter-size × fraction group. Classes were retained for visualization if the sum of their group-level mean relative abundances was ≥0.001. Classes with missing, blank, or unassigned taxonomic labels were excluded.

UpSet plots were used to compare shared and unique taxa across filter-size and filter/filtrate groups for the 16S, metagenomic, and metatranscriptomic datasets. For 16S data, taxa were agglomerated to species level using tax_glom() and converted to relative abundance. For GOTTCHA2 metagenomic and metatranscriptomic data, species-level relative abundances were summed, and group-level mean abundances were used to determine presence. For all datasets, species were considered present in a group when mean relative abundance was ≥0.001; taxa absent from all groups were removed before calculating intersections among groups.

### De novo assembly and metagenomic characterization

Quality-filtered metagenomic and metatranscriptomic reads were assembled using the MEGAHIT [v1.2.9]. To identify viral contigs, the assembled contigs were processed using geNomad [v1.9.0] and then further assessed for quality and completeness using CheckV [v0.7.0]. Only viral genomes medium, high, or complete quality were retained. For the recovery of bacterial MAGs, genome binning was restricted to the metagenomic assemblies. Specifically, metagenomic contigs were clustered into MAGs using MetaBAT [v2.12.1]. The quality of the recovered MAGs was then evaluated using CheckM [v1.2.2] to estimate completeness and contamination levels, and only MAGs medium, high, or complete quality were retained. To ensure high genomic fidelity and eliminate chimeras, GUNC [v1.0.5] was employed to flag chimeric MAGs; any MAGs identified as chimeric were strictly removed prior to all downstream taxonomic and functional analyses. All resulting MAGs were then assigned taxonomy using GTDB [v2.3.0].

## Data Availability

Raw reads are deposited on NCBI Sequence Read Archive under PRJNA1519923.

## Acknowledgements

We would like to thank Cheryl Gleaser and Erin Box for sequencing all samples. This work was funded by the Laboratory Directed Research and Development grants 202205855ECR to

M.S and 20240674ER to M.S and P.C.

## Supplementary Figures

**Supplementary Figure 1: Nucleic acid concentrations and recovered sequence read counts across wastewater fractions.** (A) Concentrations of extracted DNA and RNA derived from filter, and their corresponding filtrate fractions used as input for amplicon, metagenomic, and metatranscriptomic sequencing. (B) Total number of raw reads generated for each fraction.

**Supplementary Figure 2: Fold differences in taxonomic relative abundance between filter and filtrate fractions.** Bars show fold differences (filtrate over filter) in mean relative abundance between filtrates and their corresponding filter-retained fractions for (A) bacterial classes profiled by 16S rRNA gene amplicon sequencing, (B) fungal classes profiled by ITS amplicon sequencing, and (C) bacterial classes profiled by shotgun metagenomic sequencing.

**Supplementary Figure 3: Intersections of species-level detections across wastewater filtration fractions.** UpSet plots summarize species-level assignments shared among or detected exclusively within filter-retained material and corresponding filtrates across filtration conditions. Panels show (A) bacterial taxa profiled by 16S rRNA gene amplicon sequencing, (B) fungal taxa profiled by ITS amplicon sequencing, and (C) DNA viral taxa profiled by shotgun metagenomic sequencing. Vertical bars indicate the number of taxa in each intersection, connected dots identify the filtration conditions included in that intersection, and horizontal bars indicate the total number of taxa detected in each condition.

**Supplementary Figure 4: Mean relative abundance of bacterial classes across wastewater filtration fractions.** Heatmaps show the mean relative abundance, expressed as a proportion, of bacterial classes detected in filter and corresponding filtrates. Panels show bacterial communities profiled by (A) 16S rRNA gene amplicon sequencing and (B) shotgun metagenomic sequencing. Relative abundance for metagenomes was obtained from GOTTCHA2 taxonomic profiles. Only bacterial classes for which the summed relative abundance across all samples exceeded 0.001 are shown. Each column represents the mean across three independent filtration replicates for the indicated condition. Black cells indicate a relative abundance of zero.

## References

1. Javornik Cregeen, S., et al., *Sequencing-Based Detection of Measles in Wastewater: Texas,* January *2025*. Am J Public Health, 2025. 115(7): p. 1115–1119.

2. Galagoda, R., et al., Quantitative 16S rRNA Gene Amplicon Sequencing for Comprehensive Pathogenic Bacterial Tracking in a Municipal Wastewater Treatment Plant. ACS ES&T Water, 2023. 3(4): p. 923–933.

3. Child, H.T., et al., Comparison of metagenomic and targeted methods for sequencing human pathogenic viruses from wastewater. mBio, 2023. 14(6): p. e0146823.

4. Jiang, M., et al., Evaluation of the Impact of Concentration and Extraction Methods on the Targeted Sequencing of Human Viruses from Wastewater. Environ Sci Technol, 2024. 58(19): p. 8239–8250.

5. Pecson, B.M., et al., Reproducibility and sensitivity of 36 methods to quantify the SARS-CoV-2 genetic signal in raw wastewater: findings from an interlaboratory methods evaluation in the U.S. Environ Sci (Camb), 2021. 7: p. 504–520.

6. Gregori, I., et al., Filtering Microbial Diversity: Evaluation of Different Pore Sizes for eDNA Community Profiling. Environmental DNA, 2025. 7(6): p. e70221.

7. Djurhuus, A., et al., Evaluation of Filtration and DNA Extraction Methods for Environmental DNA Biodiversity Assessments across Multiple Trophic Levels. Frontiers in Marine Science, 2017. Volume 4 - 2017.

8. Wolfe, M.K., et al., Prospective metagenomic sequencing of wastewater across the United States yields robust viral enrichment and concordance with digital PCR measurements. medRxiv, 2026: p. 2026.05.07.26352651.

9. Liang, Z. and A. Keeley, Filtration recovery of extracellular DNA from environmental water samples. Environ Sci Technol, 2013. 47(16): p. 9324–31.

10. Tessler, M., et al., Large-scale differences in microbial biodiversity discovery between 16S amplicon and shotgun sequencing. Sci Rep, 2017. 7(1): p. 6589.

11. Kagemasa, S., et al., Metabolic Potential of Candidatus Saccharimonadia Including Rare Lineages in Activated Sludge. Environ Microbiol Rep, 2025. 17(6): p. e70231.

12. Ibrahim, A., et al., Adapted Protocol for Saccharibacteria Cocultivation: Two New Members Join the Club of Candidate Phyla Radiation. Microbiol Spectr, 2021. 9(3): p. e0106921.

13. Padilla, C.C., et al., Standard filtration practices may significantly distort planktonic microbial diversity estimates. Front Microbiol, 2015. 6: p. 547.

14. Pascoal, F., et al., Inter-comparison of marine microbiome sampling protocols. ISME Commun, 2023. 3(1): p. 84.

15. Luef, B., et al., Diverse uncultivated ultra-small bacterial cells in groundwater. Nat Commun, 2015. 6: p. 6372.

16. Murugkar, P.P., et al., Isolation and cultivation of candidate phyla radiation Saccharibacteria (TM7) bacteria in coculture with bacterial hosts. J Oral Microbiol, 2020. 12(1): p. 1814666.

17. Brinkman, N.E., et al., Reducing inherent biases introduced during DNA viral metagenome analyses of municipal wastewater. PLoS One, 2018. 13(4): p. e0195350.

18. McLaren, M.R., A.D. Willis, and B.J. Callahan, Consistent and correctable bias in metagenomic sequencing experiments. Elife, 2019. 8.

19. Rocha, U., et al., Simulation of 69 microbial communities indicates sequencing depth and false positives are major drivers of bias in prokaryotic metagenome-assembled genome recovery. PLoS Comput Biol, 2024. 20(10): p. e1012530.

20. Caporaso, J.G., et al., QIIME allows analysis of high-throughput community sequencing data. Nat. Methods, 2010. 7: p. 335.

21. Li, P.E., et al., Enabling the democratization of the genomics revolution with a fully integrated web-based bioinformatics platform. Nucleic Acids Res, 2017. 45(1): p. 67–80.

22. DeSantis, T.Z., et al., Greengenes, a chimera-checked 16S rRNA gene database and workbench compatible with ARB. Appl Environ Microbiol, 2006. 72(7): p. 5069–72.

23. Abarenkov, K., et al., The UNITE database for molecular identification and taxonomic communication of fungi and other eukaryotes: sequences, taxa and classifications reconsidered. Nucleic Acids Res, 2024. 52(D1): p. D791–D797.

24. Lo, C.C. and P.S. Chain, Rapid evaluation and quality control of next generation sequencing data with FaQCs. BMC Bioinformatics, 2014. 15: p. 366.

25. Freitas, T.A., et al., Accurate read-based metagenome characterization using a hierarchical suite of unique signatures. Nucleic Acids Res, 2015. 43(10): p. e69.

