## Supplementary material for "Characterizing the Effects of Filtration on Microbial Diversity, Reproducibility, and Genome Recovery from Wastewater": supplemetntary figures

1a)

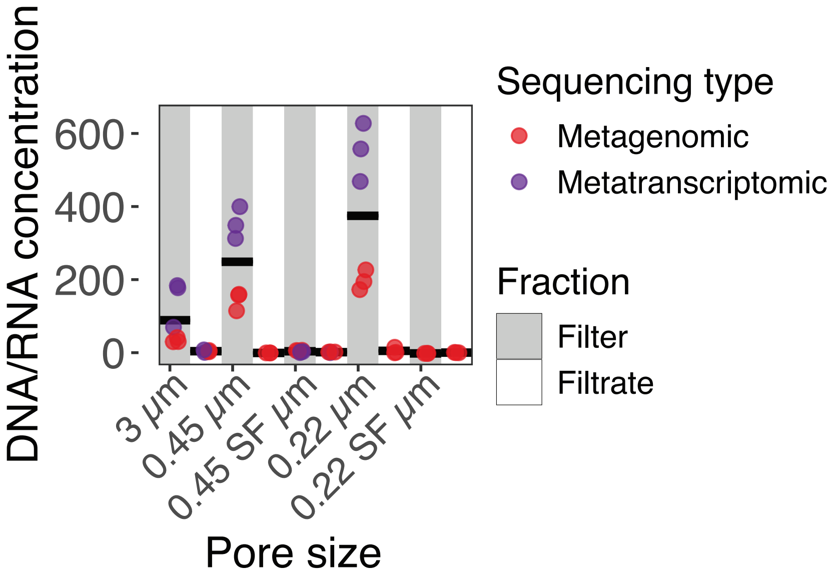
DNA/RNA concentration
600
400
200
0
3 μm
0.45 μm
0.45 SF μm
0.22 μm
0.22 SF μm
Pore size
Sequencing type
Metagenomic
Metatranscriptomic
Fraction
Filter
Filtrate

1b)

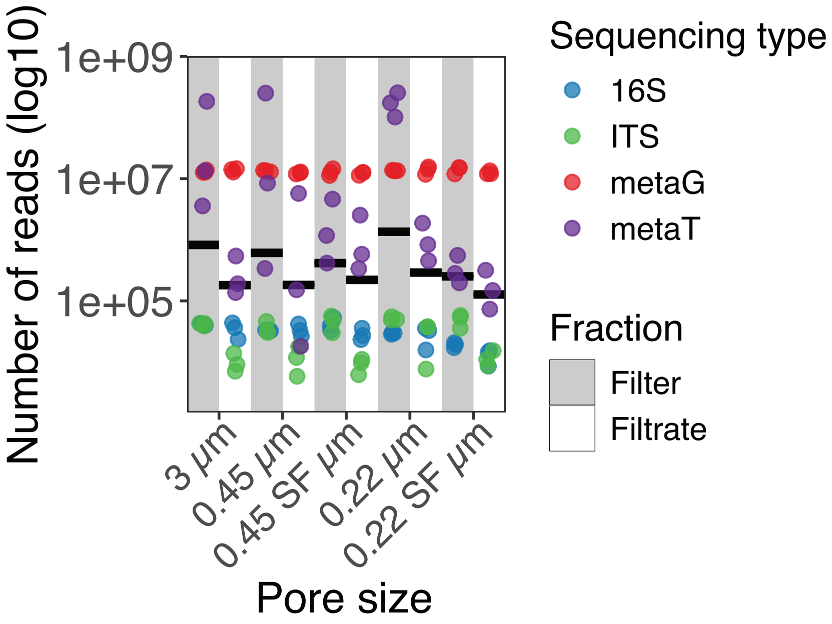
Number of reads (log10)
1e+09
1e+07
1e+05
3 μm
0.45 μm
0.45 SF μm
0.22 μm
0.22 SF μm
Pore size
Sequencing type
16S
ITS
metaG
metaT
Fraction
Filter
Filtrate

2a)

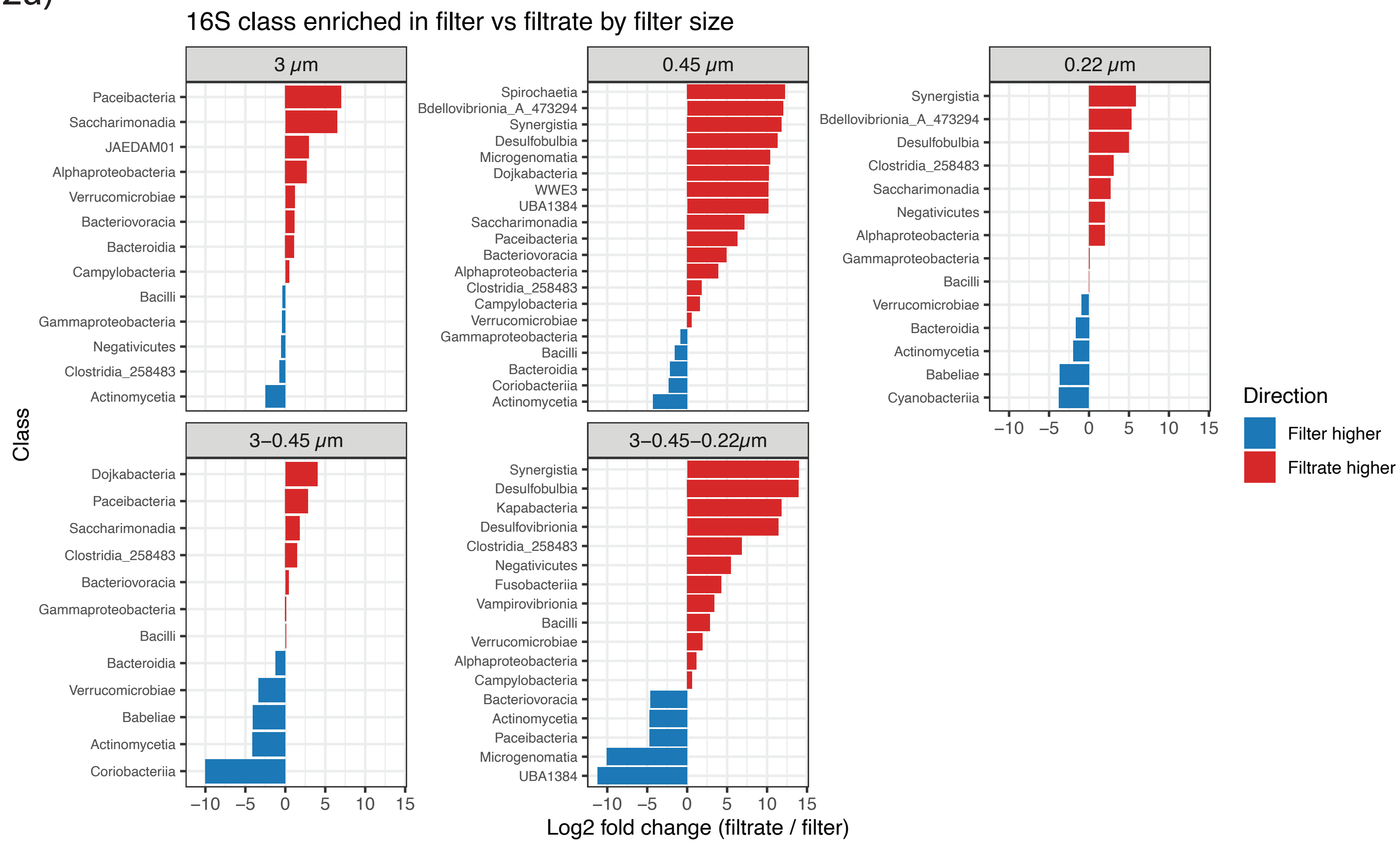
16S class enriched in filter vs filtrate by filter size
3 μm
Paceibacteria
Saccharimonadia
JAEDAM01
Alphaproteobacteria
Verrucomicrobiae
Bacteriovoracia
Bacteroidia
Campylobacteria
Bacilli
Gammaproteobacteria
Negativicutes
Clostridia_258483
Actinomycetia
0.45 μm
Spirochaetia
Bdellovibrionia_A_473294
Synergistia
Desulfobulbia
Microgenomatia
Dojkabacteria
WWE3
UBA1384
Saccharimonadia
Paceibacteria
Bacteriovoracia
Alphaproteobacteria
Clostridia_258483
Campylobacteria
Verrucomicrobiae
Gammaproteobacteria
Bacilli
Bacteroidia
Coriobacteriia
Actinomycetia
0.22 μm
Synergistia
Bdellovibrionia_A_473294
Desulfobulbia
Clostridia_258483
Saccharimonadia
Negativicutes
Alphaproteobacteria
Gammaproteobacteria
Bacilli
Verrucomicrobiae
Bacteroidia
Actinomycetia
Babeliae
Cyanobacteriia
3–0.45 μm
Dojkabacteria
Paceibacteria
Saccharimonadia
Clostridia_258483
Bacteriovoracia
Gammaproteobacteria
Bacilli
Bacteroidia
Verrucomicrobiae
Babeliae
Actinomycetia
Coriobacteriia
3–0.45–0.22μm
Synergistia
Desulfobulbia
Kapabacteria
Desulfovibrionia
Clostridia_258483
Negativicutes
Fusobacteriia
Vampirovibrionia
Bacilli
Verrucomicrobiae
Alphaproteobacteria
Campylobacteria
Bacteriovoracia
Actinomycetia
Paceibacteria
Microgenomatia
UBA1384
Class
−10 −5 0 5 10 15
Log2 fold change (filtrate / filter)
Direction
Filter higher
Filtrate higher

2b)

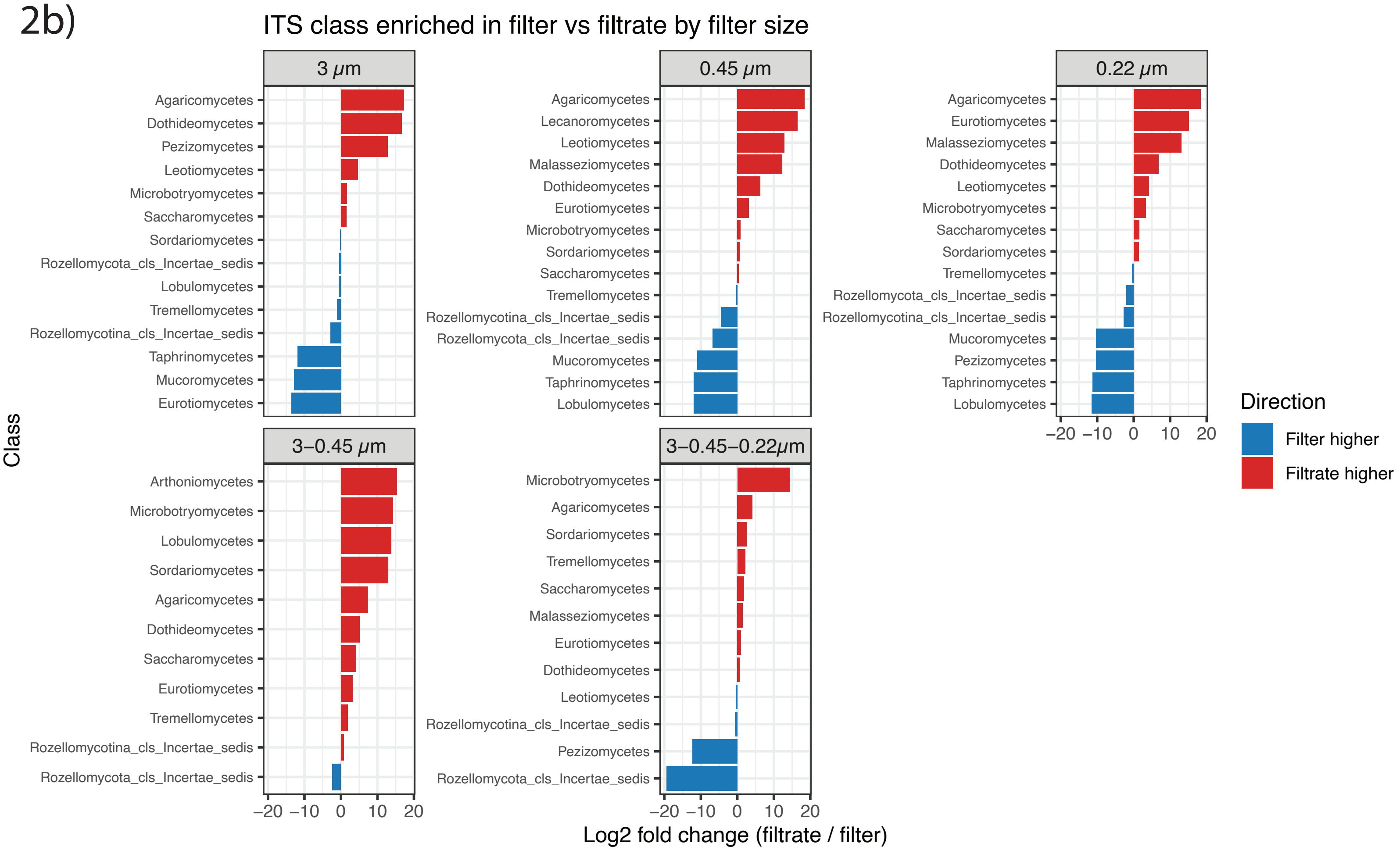
ITS class enriched in filter vs filtrate by filter size
3 μm
Agaricomycetes
Dothideomycetes
Pezizomycetes
Leotiomycetes
Microbotryomycetes
Saccharomycetes
Sordariomycetes
Rozellomycota_cls_Incertae_sedis
Lobulomycetes
Tremellomycetes
Rozellomycotina_cls_Incertae_sedis
Taphrinomycetes
Mucoromycetes
Eurotiomycetes
0.45 μm
Agaricomycetes
Lecanoromycetes
Leotiomycetes
Malasseziomycetes
Dothideomycetes
Eurotiomycetes
Microbotryomycetes
Sordariomycetes
Saccharomycetes
Tremellomycetes
Rozellomycotina_cls_Incertae_sedis
Rozellomycota_cls_Incertae_sedis
Mucoromycetes
Taphrinomycetes
Lobulomycetes
0.22 μm
Agaricomycetes
Eurotiomycetes
Malasseziomycetes
Dothideomycetes
Leotiomycetes
Microbotryomycetes
Saccharomycetes
Sordariomycetes
Tremellomycetes
Rozellomycota_cls_Incertae_sedis
Rozellomycotina_cls_Incertae_sedis
Mucoromycetes
Pezizomycetes
Taphrinomycetes
Lobulomycetes
3–0.45 μm
Arthoniomycetes
Microbotryomycetes
Lobulomycetes
Sordariomycetes
Agaricomycetes
Dothideomycetes
Saccharomycetes
Eurotiomycetes
Tremellomycetes
Rozellomycotina_cls_Incertae_sedis
Rozellomycota_cls_Incertae_sedis
3–0.45–0.22μm
Microbotryomycetes
Agaricomycetes
Sordariomycetes
Tremellomycetes
Saccharomycetes
Malasseziomycetes
Eurotiomycetes
Dothideomycetes
Leotiomycetes
Rozellomycotina_cls_Incertae_sedis
Pezizomycetes
Rozellomycota_cls_Incertae_sedis
Class
−20 −10 0 10 20
Log2 fold change (filtrate / filter)
Direction
Filter higher
Filtrate higher

2c)

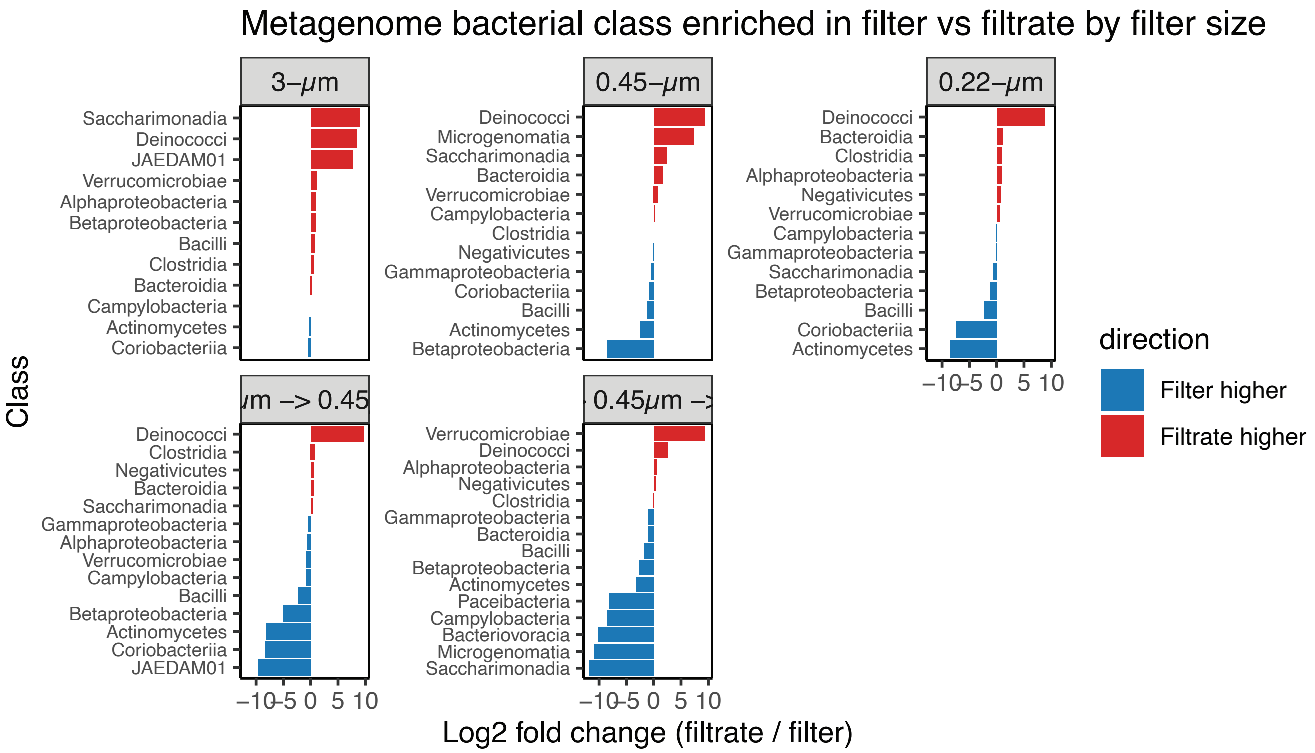

Metagenome bacterial class enriched in filter vs filtrate by filter size
3–μm
0.45–μm
0.22–μm
μm –> 0.45
0.45μm –>
Class
Saccharimonadia
Deinococci
JAEDAM01
Verrucomicrobiae
Alphaproteobacteria
Betaproteobacteria
Bacilli
Clostridia
Bacteroidia
Campylobacteria
Actinomycetes
Coriobacteriia
Microgenomatia
Negativicutes
Gammaproteobacteria
Paceibacteria
Bacteriovoracia
–10 –5 0 5 10
Log2 fold change (filtrate / filter)
direction
Filter higher
Filtrate higher


3a)

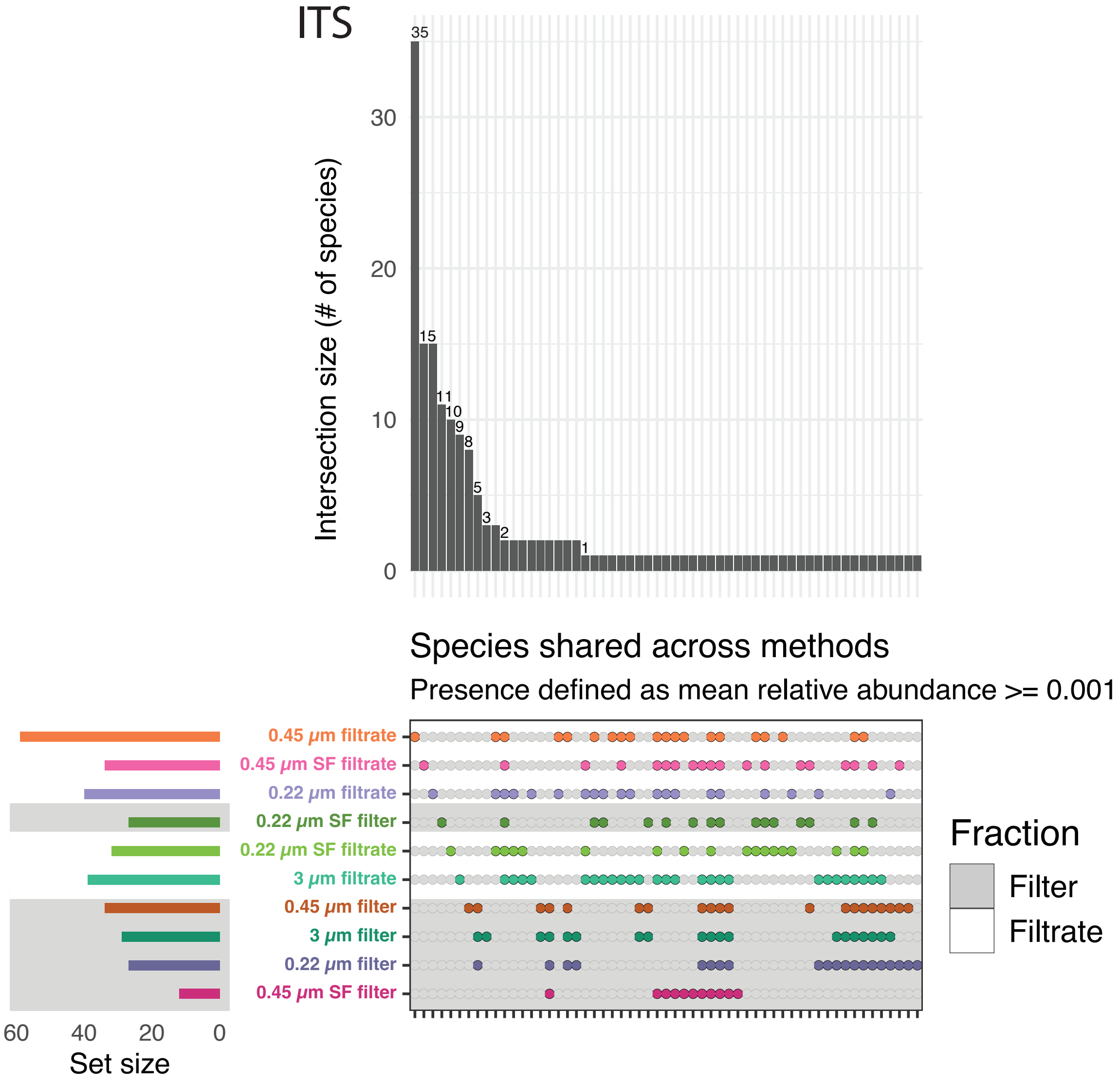

ITS
Intersection size (# of species)
35
15
11
10
9
8
5
3
2
1
0
10
20
30
Species shared across methods
Presence defined as mean relative abundance >= 0.001
0.45 μm filtrate
0.45 μm SF filtrate
0.22 μm filtrate
0.22 μm SF filter
0.22 μm SF filtrate
3 μm filtrate
0.45 μm filter
3 μm filter
0.22 μm filter
0.45 μm SF filter
60 40 20 0
Set size
Fraction
Filter
Filtrate

3b)

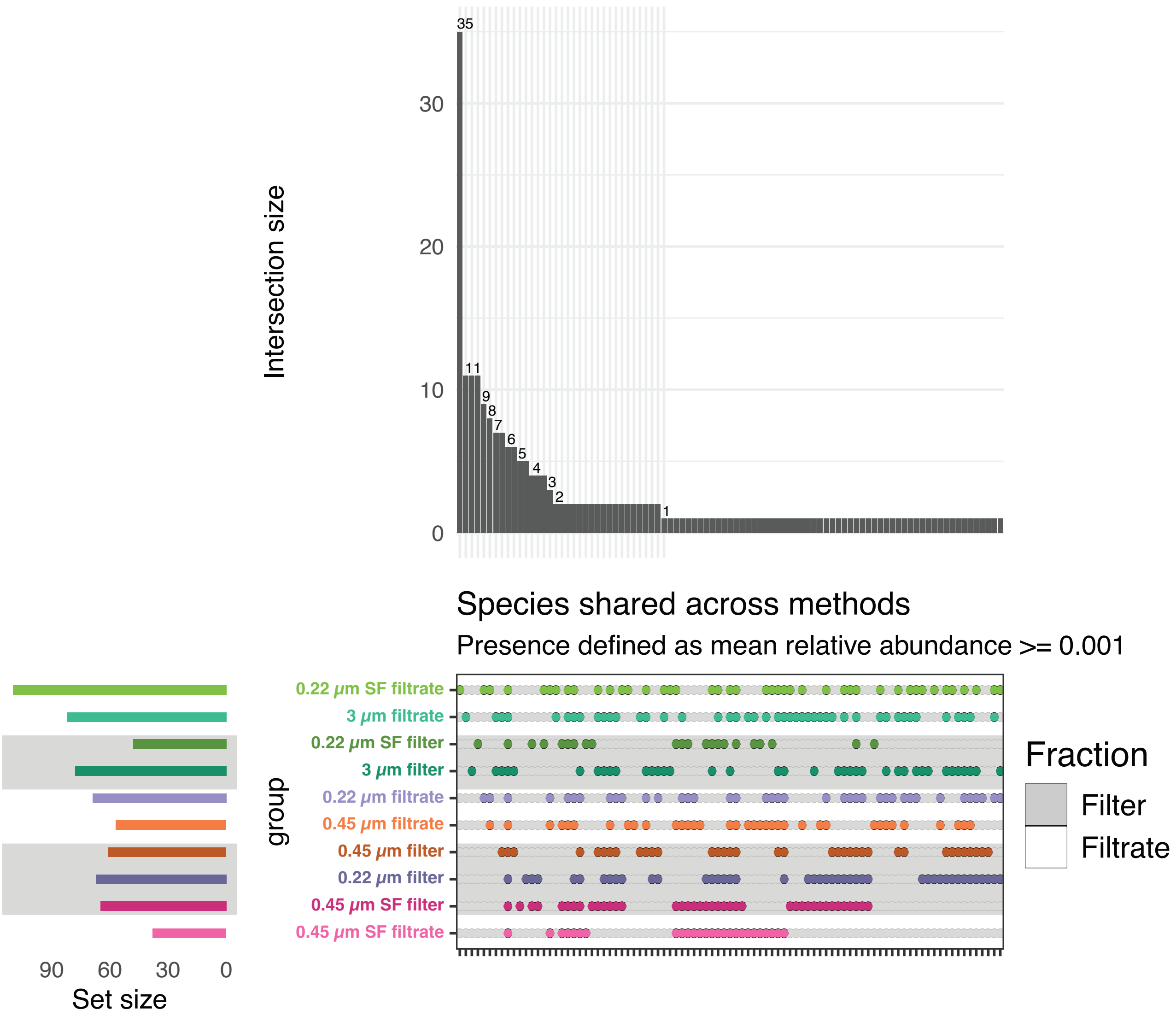
Intersection size
35
11
9
8
7
6
5
4
3
2
1
30
20
10
0
Species shared across methods
Presence defined as mean relative abundance >= 0.001
0.22 µm SF filtrate
3 µm filtrate
0.22 µm SF filter
3 µm filter
0.22 µm filtrate
0.45 µm filtrate
0.45 µm filter
0.22 µm filter
0.45 µm SF filter
0.45 µm SF filtrate
group
90
60
30
0
Set size
Fraction
Filter
Filtrate

3c)

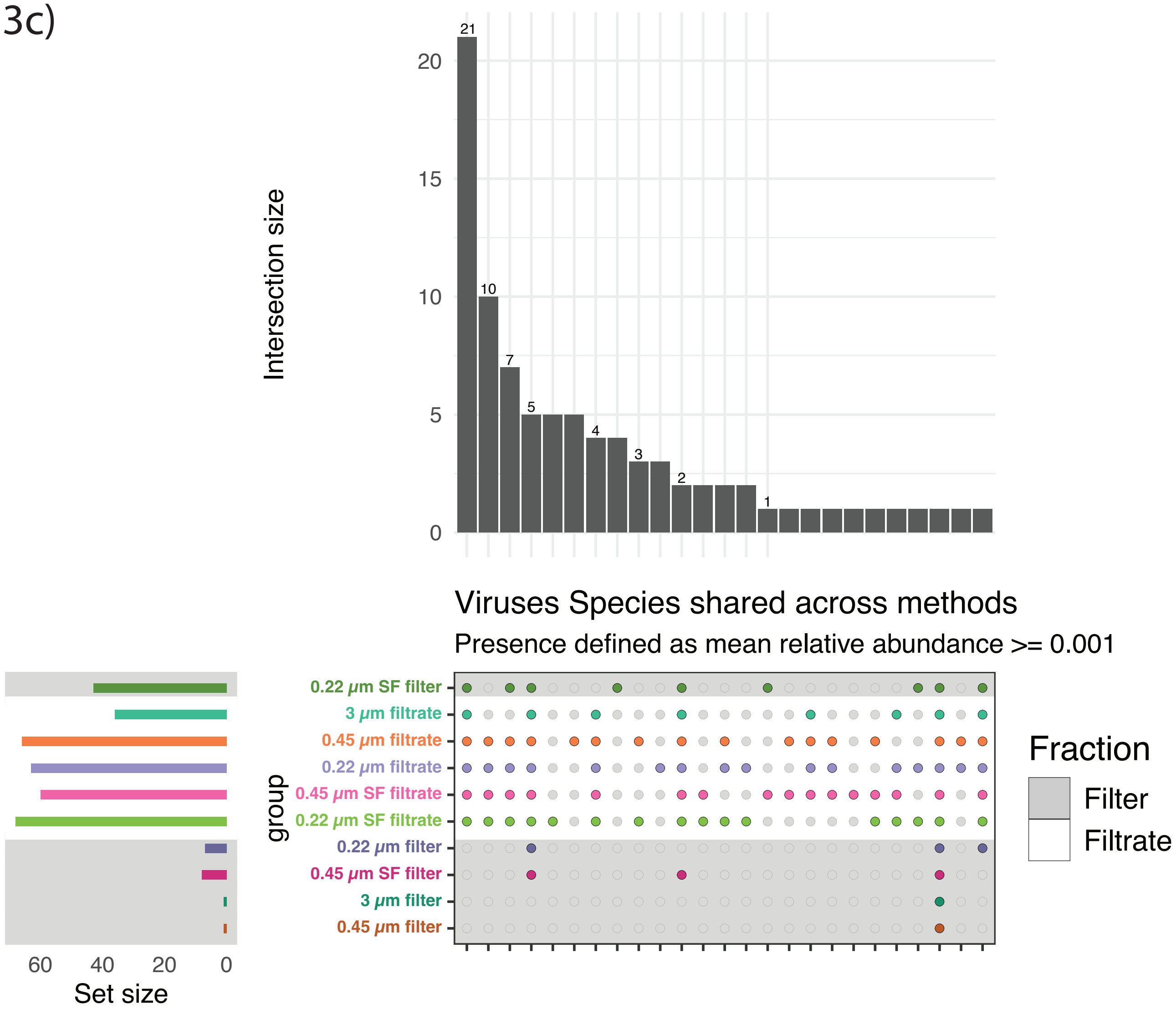
Intersection size
21
10
7
5
4
3
2
1
Viruses Species shared across methods
Presence defined as mean relative abundance >= 0.001
0.22 μm SF filter
3 μm filtrate
0.45 μm filtrate
0.22 μm filtrate
0.45 μm SF filtrate
0.22 μm SF filtrate
0.22 μm filter
0.45 μm SF filter
3 μm filter
0.45 μm filter
group
Set size
Fraction
Filter
Filtrate

4a)

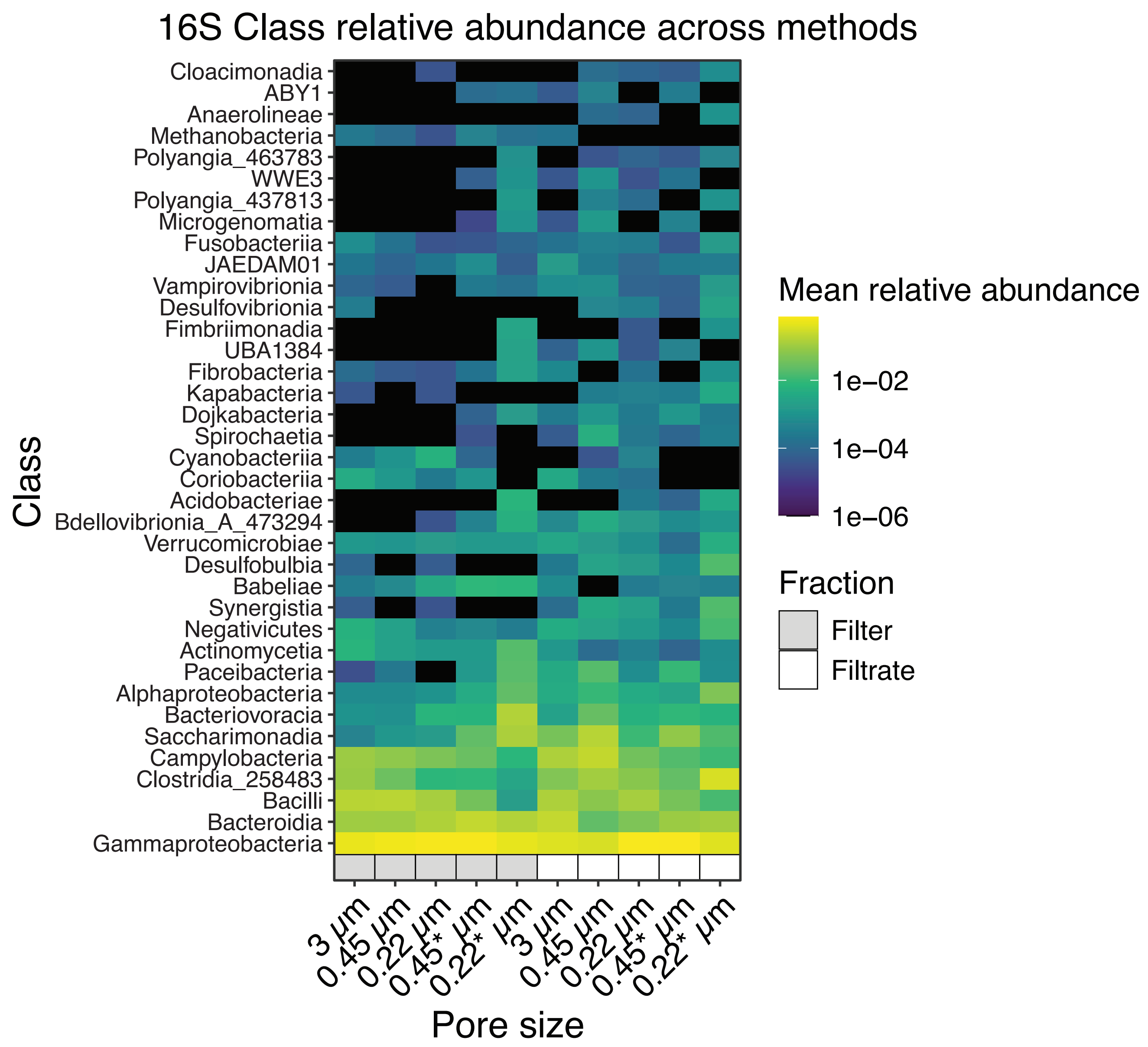
16S Class relative abundance across methods
Cloacimonadia
ABY1
Anaerolineae
Methanobacteria
Polyangia_463783
WWE3
Polyangia_437813
Microgenomatia
Fusobacteriia
JAEDAM01
Vampirovibrionia
Desulfovibrionia
Fimbriimonadia
UBA1384
Fibrobacteria
Kapabacteria
Dojkabacteria
Spirochaetia
Cyanobacteriia
Coriobacteriia
Acidobacteriae
Bdellovibrionia_A_473294
Verrucomicrobiae
Desulfobulbia
Babeliae
Synergistia
Negativicutes
Actinomycetia
Paceibacteria
Alphaproteobacteria
Bacteriovoracia
Saccharimonadia
Campylobacteria
Clostridia_258483
Bacilli
Bacteroidia
Gammaproteobacteria
Class
3 µm
0.45 µm
0.22 µm
0.45* µm
0.22* µm
3 µm
0.45 µm
0.22 µm
0.45* µm
0.22* µm
Pore size
Mean relative abundance
1e−02
1e−04
1e−06
Fraction
Filter
Filtrate

4b)

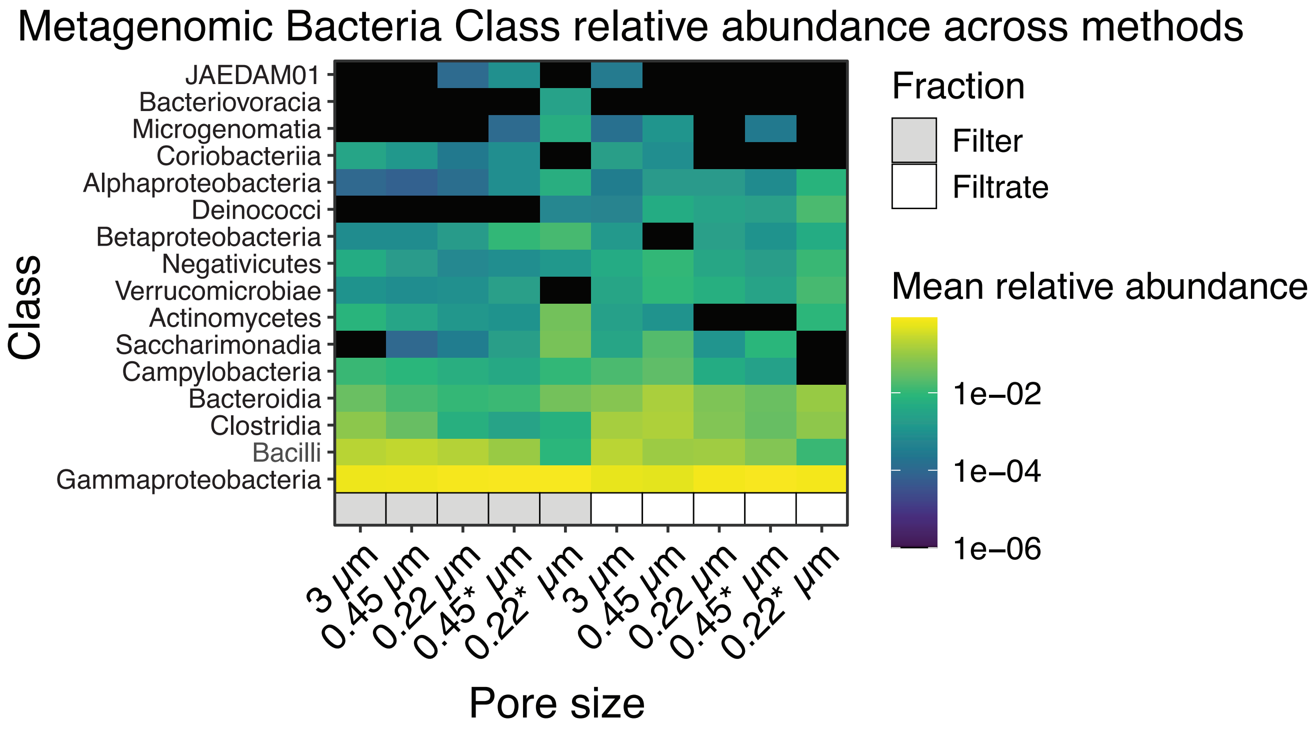
Metagenomic Bacteria Class relative abundance across methods
JAEDAM01
Bacteriovoracia
Microgenomatia
Coriobacteriia
Alphaproteobacteria
Deinococci
Betaproteobacteria
Negativicutes
Verrucomicrobiae
Actinomycetes
Saccharimonadia
Campylobacteria
Bacteroidia
Clostridia
Bacilli
Gammaproteobacteria
Class
3 µm
0.45 µm
0.22 µm
0.45* µm
0.22* µm
3 µm
0.45 µm
0.22 µm
0.45* µm
0.22* µm
Pore size
Fraction
Filter
Filtrate
Mean relative abundance
1e−02
1e−04
1e−06
